# *Placodiscus bijugus* (Sapindaceae) a new species of Endangered lowland evergreen forest tree from the western foothills of Mt Cameroon and from Korup National Park, Cameroon

**DOI:** 10.1101/2024.08.14.607904

**Authors:** George Gosline, Nouhou Ndam, Stuart Cable, Martin Cheek

## Abstract

A new range-restricted species of *Placodiscus* Radlk. is described, mapped and illustrated. *Placodiscus bijugus* is a cauliflorous tree (4 –)5 – 10 m tall, characterised by having two pairs of leaflets, 1(– 2)-flowered cymules with minute bracts, a glabrous disk, pedicels 3 – 3.2 mm long and large (3 – 4 cm diam.), shortly stipitate, retuse fruits, orbicular in outline. It is similar to *P. caudatus* Pierre ex Pellegr. which has been reported from surveys in Cameroon. We discuss the typification and range of the latter species which in fact appears to be absent from Cameroon.

*Placodiscus bijugus* is provisionally assessed as Endangered (EN B1ab(iii)+2ab(iii)) using the IUCN standard, since only three locations, with threats are reported. The species has a small, slightly range, disjunct between Mt Cameroon and Korup National Park in coastal SW region Cameroon. We review other species with this disjunct range.

## Introduction

In this paper we describe a new species of *Placodiscus* Radlk. from the lower slopes of Mt Cameroon and the Korup National Park. The species was first identified by SC and came to light in surveys for conservation management on and around the mountain (Cheek & Hepper 1994; Cheek *et al*. 1996) and was published without description as *Placodiscus* sp nov. (Cable & Cheek 1998: 127) based on five of the specimens cited here. Subsequently additional specimens were found to the west in or near the Korup National Park. The taxon is maintained in the taxonomic checklist of the vascular plants of Cameroon (Onana 2011). Here we formally publish the species so that it can be assessed for the Red List, and so that resources might be allocated for its conservation.

The genus *Placodiscus* is restricted to tropical Africa. Twenty species are accepted (POWO, continuously updated) which are distributed from Guinea-Bissau and Guinea in the West (Gosline *et al*. 2023) to the Eastern Arc Mts of Tanzania in the East (Davies & Verdcourt 1998). The genus is mainly lowland and restricted to equatorial forests, not occurring further south than DRC, and not extending to e.g. Uganda, Ethiopia or Sudan (Darbyshire *et al*. 2015). Ten species are found in West-Central Africa, or Lower Guinea sensu White (1983); seven species are confined to Africa west of the Dahomey Gap (Upper Guinea). *Placodiscus boya* Aubrév. & Pellegr. is a West African species of Ivory Coast and Ghana and records from Cameroon and Gabon are questionable (Hall 1980). Two species are found in Tanzania. None of the species are known to be adapted to savanna woodland. They are mainly shrubs and small trees, one species *P. gimbiensis* Haumann, is a liana (Fouilloy & Hallé 1973; Haumann 1960).

Morphologically, *Placodiscus* are easily recognised by the combination of their once-pinnate compound leaves without a terminal leaflet and frequentlywith prominent raised reticulate tertiary venation (in secco), the absence of petals, the unlobed disc, stamens folded in bud, and calyx tube longer than the lobes and hairy on both surfaces. In Cameroon they are distinctive for their strongly 3-lobed, fleshy but indehiscent fruit, and in flower for their complete lack of petals, the latter distinguishing them from the many species of *Chytranthus* Hook.f. which occur in the same habitat and with which they might otherwise be confused.

Represented by *Placodiscus turbinatus* Radlk. in the phylogenomic analysis of Joyce *et al*. (2023), *Placodiscus* is embedded in a largely African clade of genera where it is sister to *Haplocoelopsis africana* F.G.Davies, and these two taxa together form a subclade that is sister to a subclade consisting of *Pancovia bijuga* Willd. and *Laccodiscus ferrugineus* Radlk. Similar results had been found by Buerki *et al*. (2021), terming the 16 genera of “Clade 9” as Nephelieae Radlk. using a different delimitation to that of Radlkofer (1931). However, several African genera of Sapindaceae remain unsampled and unplaced and so the tree topology is likely to change when this shortcoming is addressed (Cheek *et al*. 2024).

The key to the species of *Placodiscus* of Fouilloy & Hallé (1973) covers all but the E African species. Because *Placodiscus* sp nov. lacks glandular hairs, has distinct petioles, has a glabrous disc, is a tree with leaves < 6 jugate, it keys to couplet 9’, where it best fits the lead to *Placodiscus boya*. However, in view of the highly raised finely reticulate venation of the abaxial leaflet surface, and the abruptly acuminate leaflet apices (caudate), together with the similar shape and dimensions of the leaflets, and the short cauliflorous spike-like inflorescences with 1-flowered cymes, the species has been confused with *Placodiscus caudatus* (*Thomas* 3319, GBIF human observations). The new species differs from both these species in the characters shown in Table 1 below.

**Table 1.** Diagnostic characters separating *Placodiscus bijugus* from *P. caudatus* and *P. boya*. Data taken from herbarium specimens at K, and also for *P. caudatus*, data from Pierre’s analysis of *Klaine* 391 and 401 (P), Fouilloy & Hallé (1973) and for *P. boya* data from Aubréville (1936) and Hall (1980).

|  | *Placodiscus caudatus* | *Placodiscus bijugus* sp.nov. | *Placodiscus boya* |
| --- | --- | --- | --- |
| Habit and height | Shrub 1(-3?) m | Tree (4-)5-10 m | Tree to 20 m |
| Leaves | 4-5-jugate | 2-jugate | 2-4-jugate |
| Pedice length at anthesis (mm) | <1 | 3-3.2 | 5-10 |
| Disc indumentum | Hairy | Glabrous | Glabrous |
| Staminal filaments (shape and indumentum) | Terete, tapering gradually from base to apex; glabrous | Dorsiventrally flattened proximally, abruptly constricted and terete distally; distally glabrous, otherwise hairy | Terete, tapering gradually from base to apex; distally glabrous, otherwise hairy |
| Fruit shape (side view) and lobe and seed number | Orbicular/3 | Orbicular/3 | Obovate/4 |
| Geographic range | Gabon and Central African Republic | South West Region, Cameroon | Ivory Coast and Ghana |

The genus is in great need of revision in the 50 years since the last major taxonomic treatment of Fouilloy & Hallé (1973) and the West African revision of Hall (1980) because a great number of additional specimens have been made, many of which appear unidentified (e.g. Sosef *et al*. 2006) and likely include several species new to science. This is the first new species to be published in the genus in the 21^st^ Century.

## Materials & methods

This study is based on the study of herbarium specimens at K and those online via gbif.org. The Flore du Cameroun volume for Sapindaceae (Fouilloy & Hallé 1973) was the principal reference work used to determine the identification of the specimens of what proved to be the new species, but the material cited in this paper was compared at the Kew herbarium (K) with specimens of all other species of the genus. All specimens cited have been seen by us unless indicated as “n.v.”. The methodology for the surveys in which most of the specimens were collected is given in Cheek & Cable (1997). Herbarium citations follow Index Herbariorum (Thiers *et al*. continuously updated), nomenclature follows Turland *et al*. (2018) and binomial authorities follow IPNI (continuously updated). Technical terms follow Beentje & Cheek (2003). The conservation assessment was made in accordance with the categories and criteria of IUCN (2012). Herbarium material was examined with a dissecting binocular microscope fitted with an eyepiece graticule measuring in units of 0.025 mm at maximum magnification.

### New Species Name

#### *Placodiscus bijugus* Gosline & Cheek *sp. nov*

Type: Cameroon, South West Region, Mount Cameroon, Onge Forest, along newly cut line made for forest inventory between mountain circular road S. of Koto II and the Onge river. Second camp of ‘transect’, proceeding W. of Masure river, 4° 21’N, 9° 02’ E, fl. 21 Oct. 1993, *Cheek* 5065 with Esowa, Riemann, Moyes (Holotype K barcode K000593390; isotypes SCA, YA).

### Description

Tree (4 –)5 – 10m tall, trunk 7 – 13.5cm diameter at 1.3 m from ground level. Bark smooth with fine fissures, light grey or brown, outer slash red brown, inner pale yellow. Branches smooth, glabrous; axillary buds c. 1mm long and wide hirsute with hairs 0.2 – 0.3mm. Leaves pinnate, 2-jugate, petiole 5.5 – 13cm long, rachis 5 – 7cm long, terete; proximal leaflets subopposite, ovate, 10 – 17 × 6 – 11cm, terminal leaflets larger 14 – 26 × 6.5 – 12.5cm, opposite, obovate; rachis terminated by conical “peg” 1 – 1.5mm long; petiolules 5 – 15 mm long by 1.5 – 2mm wide, terete, drying dark brown and wrinkled; leaflet base acuminate to attenuate, apex acuminate to spatulate 0.6 – 2.0 × 0.5 – 1.1cm, entire, with lateral nerves prominent on both sides, 6 – 9, obliquely ascending, anastomosing 2 – 5mm from margin, glabrous above and below, prominently reticulate above and below when dried, 14-16 cells per linear cm, cells 0.6 – 1.2 mm long, papery, entirely glabrous, shiny above and below. Inflorescences sparse in groups of 1 – 3 on cauliflorous bulbous projections of c. 1 cm diameter on the stem from 0.5 – 0.8 m upwards, up to the leafy stems. Inflorescences a reduced thyrse, spikes 7 – 16 cm long, rachis swollen at base, striate, puberulent with hairs < 0.1mm (appearing glabrous), flowers solitary (rarely 2), born on cymules 0.4 – 0.8 mm high and 0.5 – 0.8mm diameter, puberulent with hairs <0.2mm long, subtended by minute linear bracts 0.1 – 0.2mm long. Female flowers dull yellow, scented of powdered milk (*Cheek* 5065, K), 8.5 – 9 mm long at anthesis, including the pedicel. Pedicel terete, 3 – 3.2 × 0.8 – 0.9 mm, glabrous. Calyx campanulate, 4 – 5.5 × 3.5 – 4 mm, tube 3.5 mm long, widest c. 1mm above the pedicel apex, narrowing to 2.5 – 3.5 mm wide at the apex; calyx lobes (sepals) valvate, triangular, 1.2 – 2 × 1.5 – 1.8 mm, slightly spreading; outer surface minutely and densely hairy, hairs appressed 0.02 – 0.05mm long, pale brown; inner surface densely shortly hairy, hairs brown, simple, crisped, fine, 0.1 – 0.125 mm long. Disc shallowly saucer – shaped, 3 – 3.5 mm diam., 0.25 mm thick, glossy, black, glabrous, shallowly and irregulary lobed. Staminodes (pollen not present) 8, 3.5 – 4 mm long, included within the calyx tube; filaments 2.5 – 2.7 mm long, the proximal part dorsiventrally flattened, c. 2 mm long, 0.5 mm wide, densely white hairy, the distal part terete, c. 0.6 mm long, 0.1 – 0.13 mm wide, glabrous. Anthers basifixed, oblong, 1.1 – 1.25 × 0.6 – 0.75 mm, apex rounded, base truncate, thecae lateral, running along the margin from base to apex, glabrous. Pistil 5.5 – 6 mm long; ovary shallowly 3-lobed, 3-locular, in profile ellipsoid, 2.5 × 2.2 – 2.5 mm, completely covered in hairs; hairs pale brown, simple, straight, slightly spreading, 0.3 mm long; style triangular in transverse section, 3.25 × 0.8 – 0.9 mm, apex obtuse, the sides densely hairy, hairs dimorphic, completely covering surface, hairs simple, mainly appressed, multicellular, 0.1 mm long, apex rounded; with a few larger, spreading unicellular hairs 0.2 mm long, apex acute; stigmatic surfaces papillate, decurrent along the vertices (angled edges) for the distal 2.25 mm, 0.35 mm wide, the midline sunken, appearing as a groove. Male flowers not seen. Fruits ripening from green to yellow, 3-lobed, fleshy, indehiscent, 3- seeded; immature 1.6cm diameter, green; mature 3.5 – 4 cm long x 3.5 – 4 cm diameter, apex retuse, base with a short stipe c. 3 – 4 × 4 – 5 mm, lobes drying with transverse wrinkles, surface velvety, hairs yellow 0.1mm long. Calyx and staminal filaments persistent at base. Exocarp hard, leathery, 0.8 – 1.2mm thick. Mesocarp fleshy, scented of dried dates (*Phoenix dactylifera* L.) and prunes (*Prunus domestica* L.). Seeds dark brown-black, laterally compressed, elliptic in side view, c. 25 × 18 × 11 mm, unevenly smooth with shallow, uneven ridges, hilum not detected, dorsal edge with a groove 2 mm wide. Cotyledons superposed, white, embryo longitudinal, c. 10 × 1 mm.

**Figure 1.**
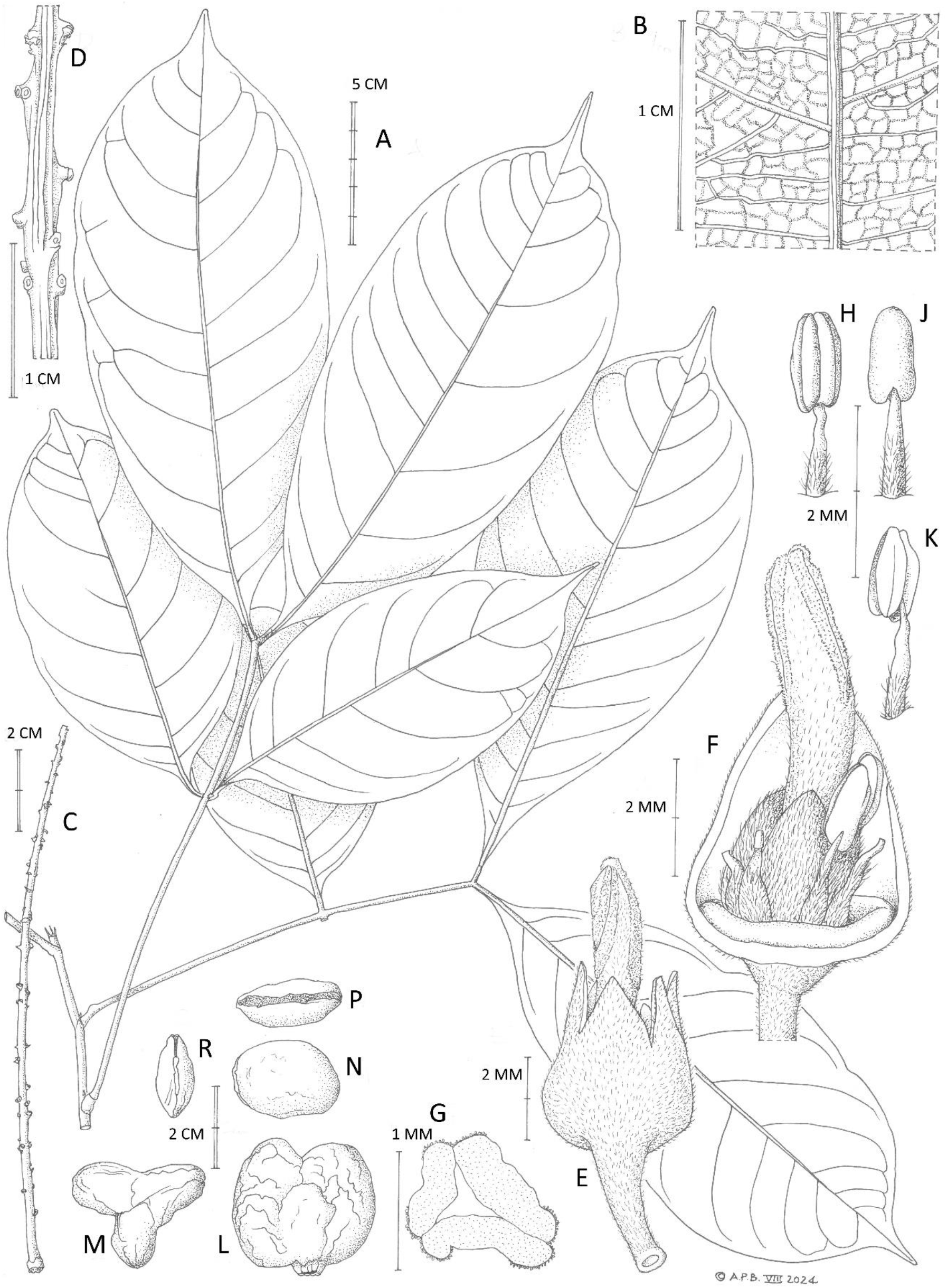
Placodiscus bijugus. **A** habit, leafy stem; **B** reticulate quaternary nervation on adaxial surface of leaflet blade; **C** axis of inflorescence (flowers fallen); **D** detail of C, showing the 1(−2)-flowered cymules; **E** female flower, side view; **F** female flower with calyx tube removed to expose disc, staminodes (several anthers removed) and pistil; **G** transverse section of the style, note the stigmatic papillate suture margins; **H, J, K** stamens emerging from the disc, respectively ventral, dorsal and side views; **L** fruit side view; **M** fruit, plan view; **N, P, R** seed, respectively side, dorsal, and end views. **A&B** from *van der Burgt* 703(K); **C-K** from *Cheek* 5065; **M-R** from *Ekema Ndumbe* 416 (K). All drawn by ANDREW BROWN.

### Recognition/Diagnosis

Differing from *Placodiscus boya* Aubrév. & Pellegr. in the leaves always 2-jugate (vs 2 – 4- jugate); inflorescences cauliflorous, inserted on the trunk within 1 m from the ground and sometimes extending to older branches (vs never cauliflorous, axillary on leafy branches to older branches); fruit orbicular in side-view, 3-lobed (vs obovate, 4-lobed).

### Distribution

Southwest Division, Cameroon, on the west slopes of Mount Cameroon and in the Korup region, two populations separated by areas of mangrove forest. We discuss the slightly disjunct distribution below. See Map 1.

**Map 1.**
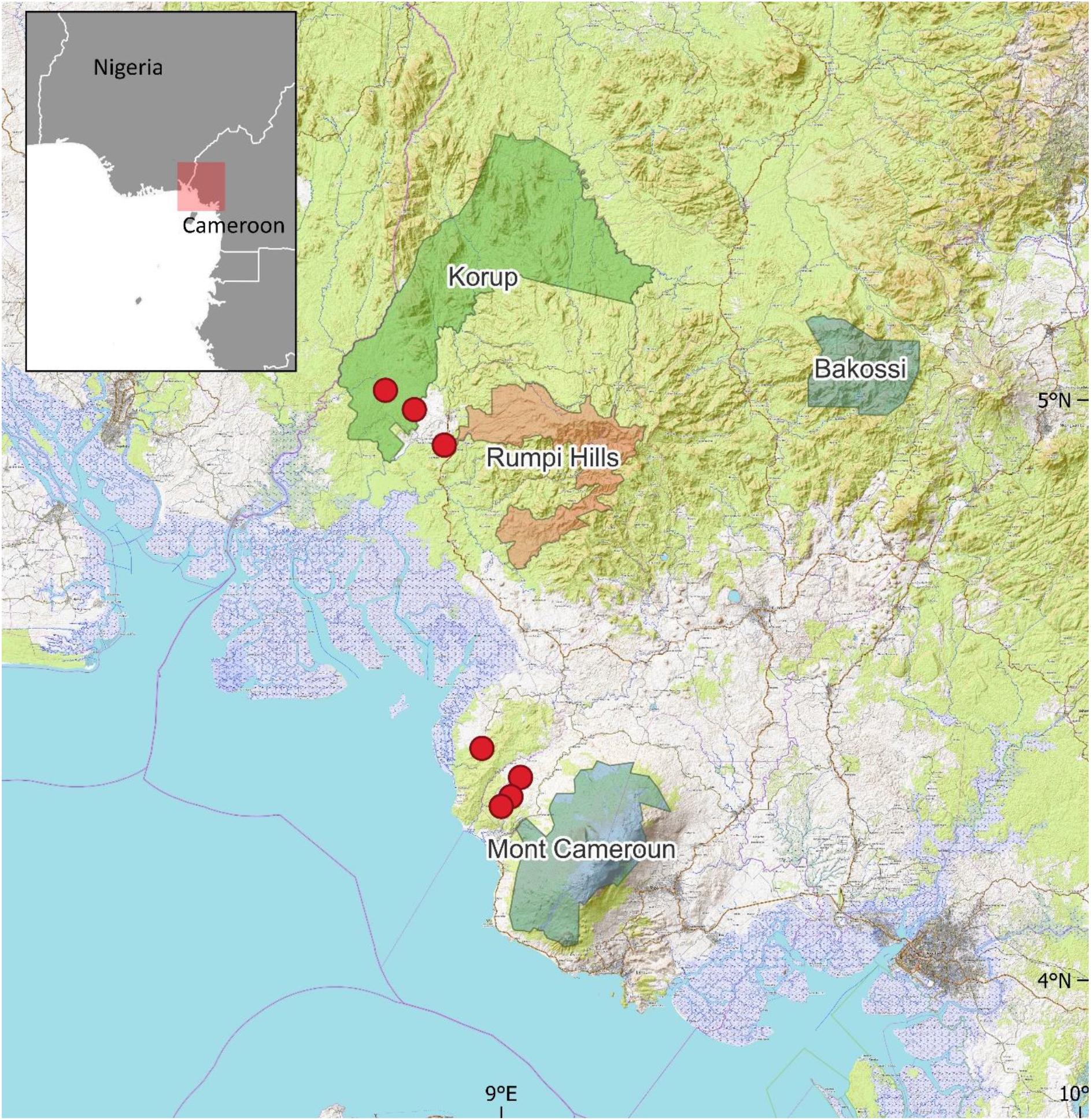
Collection sites of *Placodiscus bijugus* (red dots). Green areas = National Parks. Brown area = “Faunal reserve”.

### Habitat

Primary or secondary evergreen forest on sandy or rocky soils. On slopes or seasonally flooded sandy areas.

### Conservation status

The new species has an extent of occurrence of 525 km^2^ and area of occupancy of 28 km^2^ as calculated using GeoCAT (Bachman *et al*. 2011). There are only three “threat-based” locations (IUCN 2012), the Onge forest (western slopes of Mt Cameroon), the Korup National Park, and the Besingi community forest near Korup. The Onge forest is threatened with forest clearance, both for timber and oil palm (Cheek *et al*. 2021), as is also the forest at Besingi due to small-holder agriculture. Korup has been considered protected, but during the 2016-ongoing wars, also known as the Anglophone crisis, forest guards are reported to have fled the Korup, leaving it unprotected, and in other protected areas, armed groups or civilians seeking security, have moved in, clearing habitat (Bang 2022; Tabi *et al*. 2020). Species within the genus *Placodiscus* are slow growing trees and shrubs and are indicators of good quality, undisturbed forest. They do not regenerate well after major forest disturbance and therefore appear sensitive to the threats posed by forest clearance (Gosline *et al*. pers. obs. 1995 – 2016). It is to be hoped that the species will be found at further sites, however the prospect of this seems low since numerous intensive botanical surveys have been conducted in neighbouring areas without encountering this species (Chapman & Chapman 2001; Cheek *et al*. 2000; 2004; 2006; 2010; 2011; Harvey *et al*. 2004; 2010; Maisels *et al*. 2000). Given the threats and restricted range, an assessment of Endangered (EN B1ab(iii)+2ab(iii)) is here given to this species.

### Specimens examined

#### CAMEROON

**South West Region (formerly Province), Mount Cameroon, Onge Forest**, Bomana-Koto road c. 500 m, bearing 305° towards Onge R., 5 hr walk from the road, 4°19’N, 9°01’E, 400 m alt., fr., 20 Oct. 1993, *Ndam* 754 (SCA); Forest to the W of Onge R. about 6 km West of Liwenyi village (c 14 km North of Idenau), 4°24’N, 8°58’E, 200 m alt., fr. 11 Oct. 1993, *Watts* 873 (K, SCA n.v., YA n.v.); Onge River, secondary forest between OB2 and OB3 on transect OB. 04°15’N, 09°01’E, 400 m alt., fr., 19 Oct. 1993, *Tchouto Mbatchou* 878 (K, SCA n.v., YA n.v.); Along newly cut line made for forest inventory between mountain circular road S. of Koto II and the Onge river. Second camp of ‘transect’, proceeding W. of Masure river, 4°21’N, 9°02’E, 350 m alt., fl.,fr., 21 Oct.1993, *Cheek* 5065 (K holo.; iso. SCA, YA); Bomana, about 4.km from roadside, 04°18’N, 09°00’E, 230 m alt., fr., 08 Oct. 1993, *Akogo* 89 (K, SCA n.v.); Onge, field notes lost, fr., 08 Oct. 1993 *Ekema Ndumbe* 416 (K, SCA n.v.); **Korup area**: Besingi, near Besingi village, downstream of high bridge over Idu river 50 m alt., fr., 14 Sept. 2006, *Burgt,van der* 847 (K, MO n.v., P n.v, WAG n.v., YA n.v.); Korup National Park: P plot, subplot 9I, 05°01’N, 08°48’E, alt. 100 m, fl., fr., 31 Aug. 2004, *Burgt, van der* 703 (K, BR n.v., G n.v., MO n.v., P n.v., SCA n.v., WAG n.v., YA n.v.); Transect S “PAPI”, 04°59’N, 08°51’E, fr. Apr. 1979, *Thomas D*.*W*. 748 (K, SCA n.v.).

### Notes

#### Sexuality

Acevedo-Rodriguez *et al*. (2011) discuss the sexuality of Sapindaceae. *Placodiscus* is usually described, following Radlkofer (1931) as “falsely polygamous”. Sapindaceae are generally dichogamous (or duodichogamous) (Acevedo-Rodriguez *et al*. 2011) with male flowers preceeding female flowers, supposedly on the same inflorescence. All our specimens have female inflorescences (or infructescences); the female flowers have robust but sterile anthers (staminodes). Fruiting plants are more noticeable to collectors, and collections of functionally male flowers in the genus are almost absent at Kew.

#### *Placodiscus caudatus*, typification and identity

The name *Placodiscus caudatus* was coined by the French botanist Louis Pierre in his extensive notes made in Paris when identifying the collections that he had acquired probably by purchase, from Klaine. These collections were made in the area of Libreville, now Gabon when Klaine was a missionary. Some hundreds of new names were coined by Pierre in this way, but he only took a few forward to publication, many being taken up by other botanists after his materials were donated to P in 1904, before his death. This resulted in the authority of most of the names being credited as “ Pierre ex…”

*Placodiscus caudatus* is written on the seven pages of Pierre’s notes that are associated at P with the specimens *Klaine* 391 and *Klaine* 401. They are also accompanied by an analytical drawing of the flowers made in pencil, probably by Delpy, who worked as an artist for Pierre. Several of the many drawings he did of other species for which Pierre coined new names were completed in ink and reproduced by a form of photocopying, distributed to other herbaria and have been considered under the Code by Breteler as validating Pierre’s names (Breteler 2006). This was not the case with *Placodiscus caudatus*. Instead, the name was taken up by two other botanists. The first by Pellegrin (1924) who like Pierre was working up for publication the collections of another French collector, in this case Le Testu, from a different part of Gabon, Tchibanga, on the southern border with what is today the Republic of Congo. Pellegrin applied Pierre’s name to *Le Testu* 1627 publishing it in his Flore du Mayombe with a short description as Pierre ex Pellegr. Independently Radlkofer in his monographic work on Sapindaceae, also published the name as Pierre ex Radlk. But based explicitly on Pierre specimens (Radlkofer 1931). Subsequent works, such as the treatments of Sapindaceae for Flore du Gabon, and also Flore du Cameroun, have followed Radlkofer and generally not referred to Pellegrin (e.g. Fouilloy & Halle 1973). This may be because *Le Testu* 1627 appears not to be at P, and indeed has not been traced by us on gbif.org. Nevertheless, this at first sight appears to be the type specimen of *Placodiscus caudatus* Pierre ex Pellegr. which predates Radlkofer’s publication. Should this specimen be refound, it is possible that, being geographically distant from Klaine’s Libreville collected specimens, and with differences in the morphology indicated in the description, it is a different taxon. Until *Le Testu* 1627 is found or another specimen is made from Tchibanga that fits the description, the question will remain open. For the present we are assuming that Pellegrin, in Paris was correct in identifying *Le Testu* 1627 with Pierre’s Klaine specimens at P, and so the Klaine specimens characterise the species *Placodiscus caudatus* Pierre ex Pellegr.

The large number of human observations of *Placodiscus caudatus* in Cameroon listed on GBIF (gbif.org continuously updated) are all from a survey of the Rumpi Hills Forest Reserve which is adjacent to the area of occupation of known *P. bijugus* specimens. It is possible that these observations are in fact *P. bijugus*, or of the undescribed species mentioned below, but without voucher specimens it is not possible to be sure. In any case, it is unlikely that they are *P. caudatus* which we now doubt is present in Cameroon despite being included there in Onana & Cheek (2011) and Cheek (2004).

#### Other new species

Five other specimens from the same area appear to be another undescribed species, informally here named “*Placodiscus sp. nov*. 2”. They are superficially similar to *P. bijugus* but are not bijugate (they have 6-7 pairs of leaflets) and differ in leaf shape (bases attenuate) and number of flowers/fruits in a cymule (3 or more vs 1(−2)). We lack flowering material, so a complete description is not yet possible.

#### Species shared between southern Korup and western Mt Cameroon

Thanks to the pioneering work of Duncan Thomas (Thomas 1986; Thomas *et al*. 2003), the Korup National Park has been the site of discovery of many plant species new to science (Breteler 1990; Cheek 2002; Cheek *et al*. 2013; Kenfack *et al*. 2018; Sainge *et al*. 2005, Stone *et al*. 2008; van der Burgt 2009, 2010, 2016; van der Burgt & Newbery 2006), including the new genus and species *Korupodendron songweanum* Litt & Cheek (Litt & Cheek 2002).

Many near-endemic species (present in Korup and one or two other locations) are generally more highly threatened than those that are endemic to Korup e.g. *Glumea korupensis* Burgt and *Gambeya korupensis* Ewango & Kenfack which both also occur just outside the National Park, (van der Burgt & Newbery 2006; Ewango *et al*. 2016). This is because their sites outside Korup are usually much more highly threatened than those inside Korup. As with *Placodiscus bijugus*, a standard pattern for such near endemics is that their location outside Korup is in the western forests of Mt Cameroon 50 – 80 km to the SE, in either the Mokoko or Onge forests, e.g. *Tessmannia korupensis* Burgt (van der Burgt 2016). Like southern Korup, both Mokoko and Onge forests are also based, at least partly, on highly leached impoverished soils. In the other direction, several threatened species first discovered in these western forests of Mt Cameroon and at first seemingly endemic there, were later found in Korup e.g. *Belonophora ongensis* Cheek & S.E.Dawson (Cheek & Dawson 2000), *Cola suboppositifolia* Cheek (Cheek 2002) and *Drypetes burnleyae* (Cheek *et al*. 2021). However, the western forests of Mt Cameroon also have global endemics that have not been found in the lowland forests of Korup e.g. endemics to Mokoko such as *Octoknema mokoko* Gosline & Malécot (Gosline & Malécot 2011) and *Dracaena mokoko* Mwachala & Cheek (Mwachala & Cheek 2012).

## Conclusion

Most (three out of four) new plant species described today are already threatened with extinction (Brown *et al*. 2023), so description of new species such as that in this paper is urgent. Until species are formally described, they are invisible to science and their Red Listing is impeded (Cheek *et al*. 2020). Cameroon already has the highest documented number of extinct plant species in tropical Africa (Humphreys *et al*. 2019). Most of these extinctions have occurred in the lowland forests of Mt Cameroon due to industrial crop plantations; e.g. *Oxygyne triandra* Schltr. (Thismiaceae, Cheek *et al*. 2018) and *Afrothismia pachyantha* Schltr. (Afrothismiaceae, Cheek *et al*. 2019). If *Placodiscus bijugus* is not to follow this path to extinction, improved conservation and monitoring measures will be needed in Cameroon, e.g. Important Plant Area programmes (Darbyshire *et al*. 2017). It is helpful that both Korup and Onge-Mokoko are now Important Plant Areas (Murphy *et al*. 2023).

## Declarations

The authors declare that they have no conflict of interest.

## Acknowledgements

This paper was completed as part of the Cameroon TIPAs (Tropical Important Plant Areas) project at RBG, Kew, which is supported by Players of Peoples Postcode Lottery. Fieldwork funding in the 1990s leading to the discovery and collection of most of the specimens cited in this paper was received from the former Overseas Development Administration (ODA) of the UK government (now incorporated in the Foreign, Commonwealth and Development Office) through the former Mount Cameroon Project at Limbe Botanic Garden, which was co-managed by ODA with the Forest Department of the Cameroon Government. The fieldwork during which the type collection was collected was supported by the Earthwatch Institute with the support of volunteers of Earthwatch Europe, Oxford (Sally Moyes and Polly Riemann) and others by our colleagues including the late Martin Etuge and the late Ekema Ndumbe. The most recent collections of the new species from Korup were made possible by a visit sponsored by the Bentham-Moxon Trust of RBG, Kew. This paper is one of a series resulting from the partnership between RBG, Kew and IRAD-National Herbarium of Cameroon, Yaoundé and we thank the late Dr Benoît Satabié, Drs Gaston Achoundong, Florence Ngo Ngwe, Eric Nana, Jean Lagarde Betti, Barthelemy Tchiengué, the current and former directors or acting Directors, of IRAD-National Herbarium of Cameroon, Yaoundé, for expediting the collaboration between our two institutes.

